# Synthetic Biology Research and Innovation Profile 2018: Publications and Patents

**DOI:** 10.1101/485805

**Authors:** Philip Shapira, Seokbeom Kwon

**Affiliations:** Manchester Institute of Innovation Research, Manchester Business School, University of Manchester, Manchester, M13 9PL, UK; Manchester Synthetic Biology Research Centre for Fine and Speciality Chemicals (SYNBIOCHEM), Manchester Institute of Biotechnology, University of Manchester, Manchester, UK; School of Public Policy, Georgia Institute of Technology, Atlanta, GA 30332-0345, USA

## Abstract

A profile of synthetic biology research and innovation is presented using data on publications and patents worldwide and for the UK and selected benchmark countries. The search approach used to identify synthetic biology publications identifies a core set of synthetic biology papers, extracts and refines keywords from these core records, searches for additional papers using those keywords, and supplements with articles published in dedicated synthetic biology journals and curated synthetic biology special collections. For the period from 2000 through to mid-July 2018, 11,369 synthetic biology publication records are identified worldwide. For patents, the search approach uses the same keywords as for publications then identifies further patents using a citation-tree search algorithm. The search covered patents by priority year from 2003 to early August 2018. Following geographical matching, 8,460 synthetic biology basic patent records were identified worldwide. Using this data, analyses of publications are presented which look at the growth of synthetic biology outputs, top countries and leading organizations, international co-authoring, leading subject categories, citations, synthetic biology on the map of science, and funding sponsorship. For patents, the analysis examines growth in patenting, national variations in publications compared with patenting, leading patent assignees, and the positioning of synthetic biology on a visualized map of patents.

## Overview of Methods and Sources

In this paper, we provide an overview profile of synthetic biology research and innovation using data on publications and patents worldwide and for the UK and selected benchmark countries.

The data source for publications is the Web of Science (WOS). This is a leading large-scale database of bibliometric and citation records.^1^ The search approach used to identify synthetic biology publications within the WoS follows the procedures described in Shapira et al. (2017).^2^ The approach identifies a core set of synthetic biology papers, extracts and refines keywords from these core records, searches for additional papers using those keywords, and supplements with articles published in dedicated synthetic biology journals and curated synthetic biology special collections. The search covers synthetic biology publications recorded in WoS from 2000 through to mid-July 2018.^3^ VantagePoint textmining software is used to clean and analyze the publication records.^4^ Records were de-duplicated using the WOS ISI Unique Article Identifier. In total, the combined and cleaned data set comprises 11,369 synthetic biology publication records.

For patents, the main data source is Derwent Innovations, which offers a recognized and curated global database of patents and patent applications.^5^ The search approach used to identify synthetic biology patents follows Kwon et al. (2016).^6^ The approach employed the same keywords as for publications (see Shapira et al., 2017) to identify an initial set of patent records. Further patents were identified using a citation-tree search algorithm. The search covered patents by priority year from 2003 to 2018 (as of Aug 3, 2018), with 9,263 Derwent Innovations patent records identified. Geographical address information for assignees was obtained by record matching with PATSTAT.^7^ This identified 8,498 matching patent records (including 8,460 records with basic patent numbers for the original invention in a patent family).

As an emerging, dynamic, multidisciplinary domain, there are multiple complexities in defining synthetic biology, as well as constraints in the available data sources. For detailed discussions of these issues, the search methods adopted, and their strengths and limitations when applied to identifying synthetic biology publications and patents, reference should be made to the Shapira et al. (2017) and Kwon et al. (2016) respectively.

The following sections present key results from our analyses of synthetic biology publication and patents.

## Synthetic Biology Publications

### Global Growth of Synthetic Biology Publications

Synthetic biology publications as identified in the WoS increased, on an annualized basis, from under 200 publications worldwide in the early 2000s to over 500 in 2010. We estimate that worldwide publications will reach about 1450 in 2018. The four leading countries, by authorship, in producing synthetic biology publications are the USA, the UK, China, and Germany. (Figure 1.) Authors from these four countries account for more than 70 per cent of all synthetic biology publications from 2000 through to the present (Table 1).

**Figure 1.**
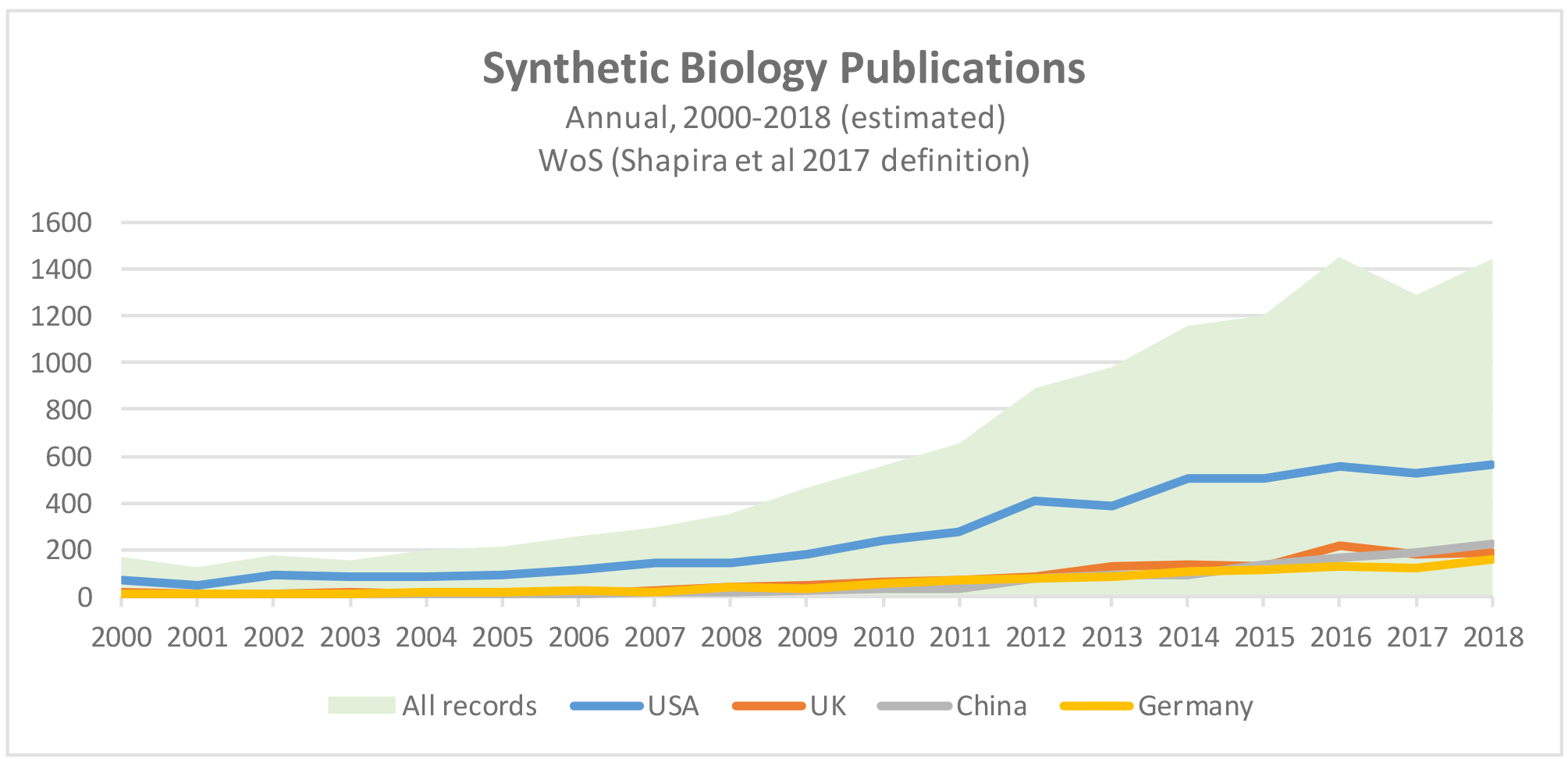
Synthetic Biology Publications. Source: Analysis of Web of Science publication records (2000 to mid-July 2018), Shapira et al. 2017 synthetic biology search strategy, N=11,369. Annualized totals for 2018 estimated from part-year 2018 publication trends.

Table 1 shows the top 15 countries for synthetic biology publications for the 2000-2017 period, based on author countries. The USA is the leading country, producing more than two-fifths of the world’s synthetic biology publications, although its share of the world total has dipped across the three periods depicted in the table. The UK, China and Germany are the next most prolific countries for synthetic biology publications over the period 2000-2017. While UK publications have grown strongly over this period, China’s have recently grown at a faster rate. In 2015 and 2017, China achieved comparable output to the UK in terms of annual synthetic publications; in 2018 (papers published to date) Chinese authors published more papers (120) than UK authors (101). Overall, authors in about 90 countries have published synthetic biology publications (Figure 2).

**Table 1.**
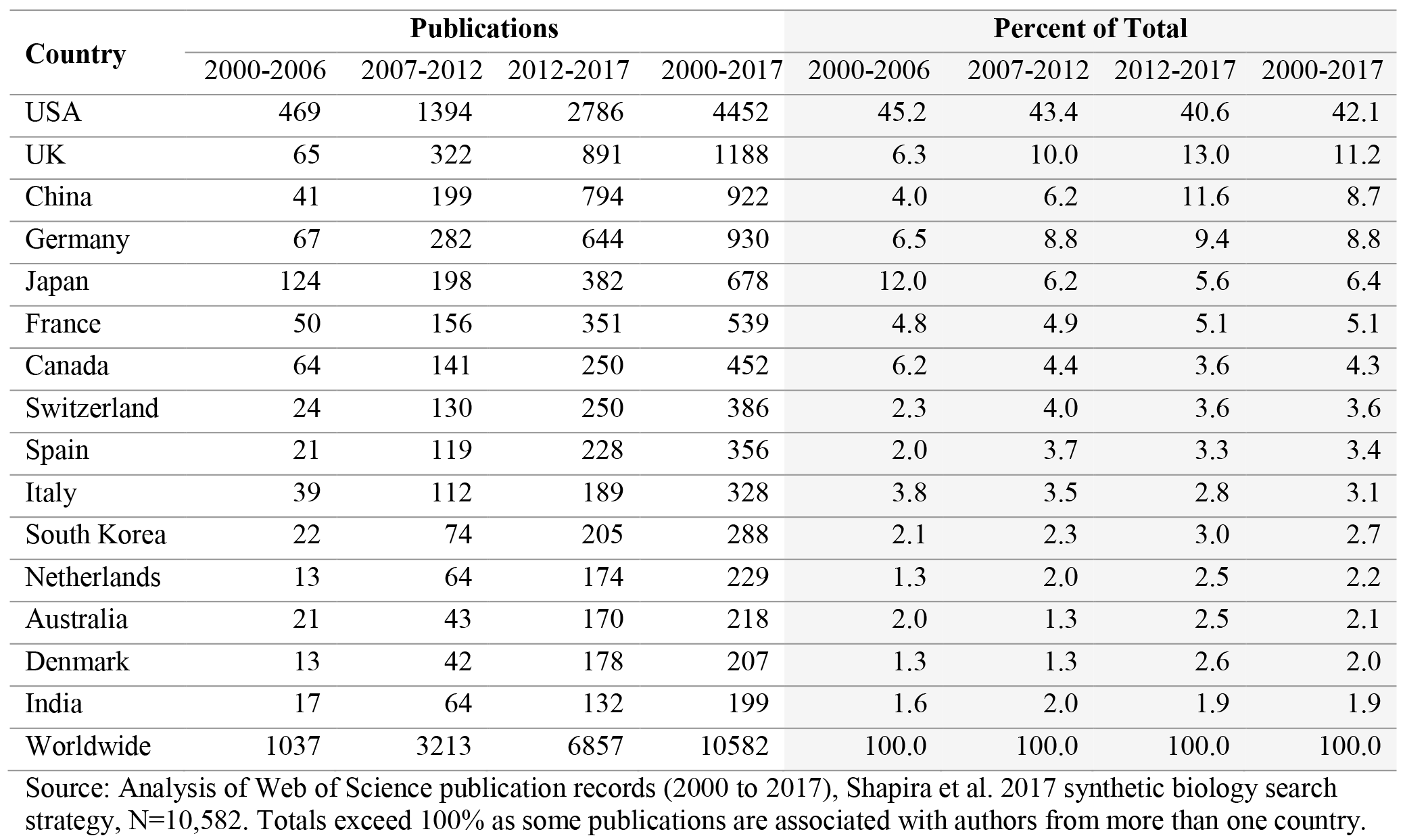
Top 15 Countries for Synthetic Biology Publications.

**Figure 2.**
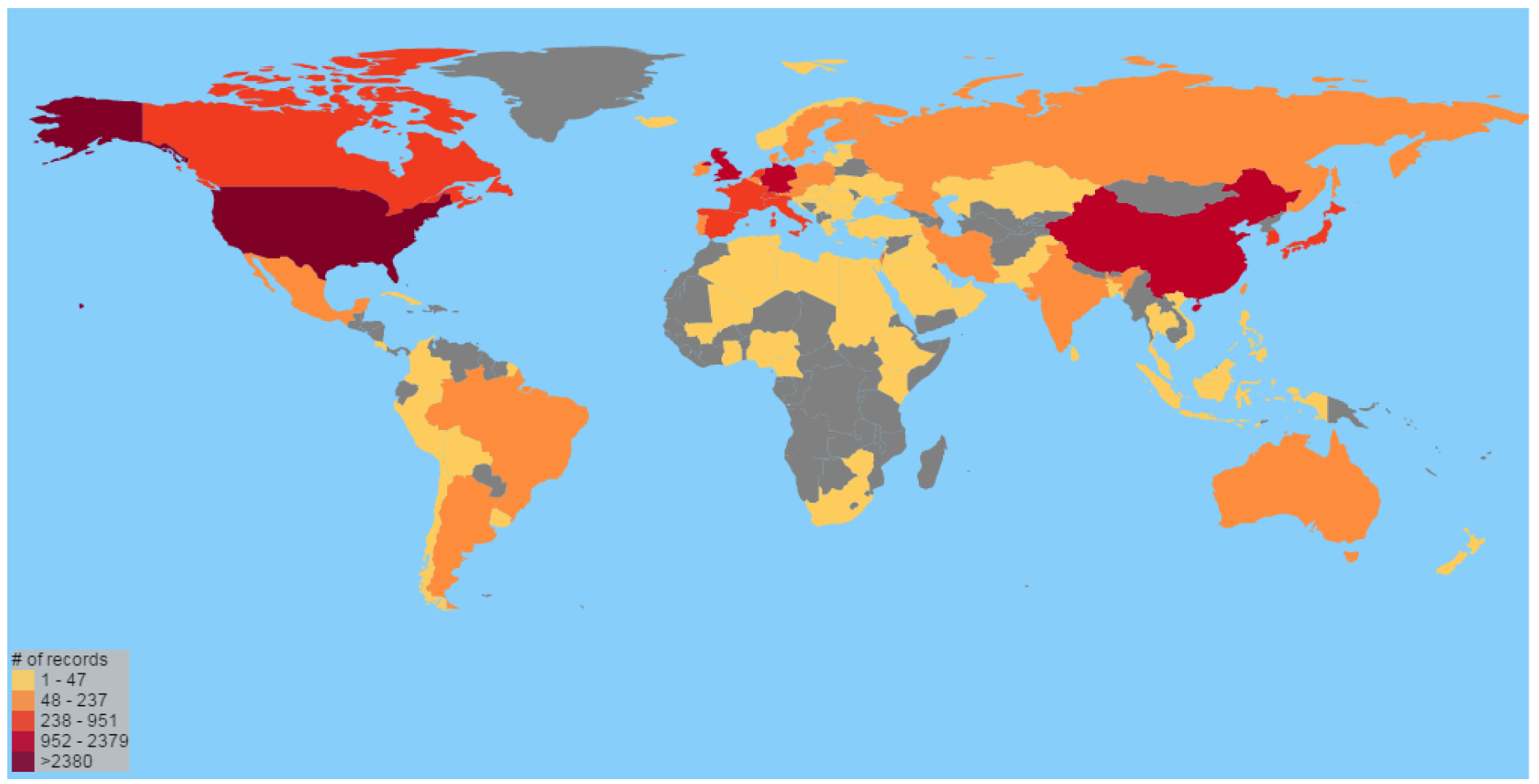
Synthetic Biology Publications by Country, 2000-2018*. Source: Analysis of Web of Science publication records (2000 to mid-July 2018), Shapira et al. 2017 synthetic biology search strategy, N=11,369. *Part Year.

### Top Organizations for Synthetic Biology Publishing

Table 2 shows the top 50 institutions (in terms of publication counts) worldwide involved in synthetic biology research publishing, by author affiliation. These top 50 institutions accounted for about one-half (49.8%) of all worldwide synthetic biology publications from 2000 through to mid-2018. Most of the top publishing institutions are universities, with representation also from other academic organizations, public research organizations, and governmental laboratories.

**Table 2.** Top 50 World-Wide Affiliations for Synthetic Biology Publications (2000-2018*)

| **Author Affiliation** | **Records** | **Author Affiliation** | **Records** |
| --- | --- | --- | --- |
| **1** MIT | 330 | **26** Osaka Univ | 91 |
| **2** Harvard Univ | 278 | **27** Univ Calif Davis | 89 |
| **3** Univ Calif Berkeley | 274 | **28** Joint BioEnergy Inst | 88 |
| **4** Swiss Fed Inst Technol | 222 | **29** CSIC | 85 |
| **5** Chinese Acad Sci | 211 | **30** Cornell Univ | 84 |
| **6** Imperial Coll London | 185 | **31** Univ Cambridge | 84 |
| **7** Univ Illinois | 155 | **32** CNRS | 83 |
| **8** Univ Toronto | 152 | **33** Japan Sci & Technol Agcy | 81 |
| **9** Stanford Univ | 149 | **34** Univ Basel | 81 |
| **10** Univ Calif San Diego | 136 | **35** Univ Texas Austin | 77 |
| **11** Univ Tokyo | 134 | **36** Univ Wisconsin | 77 |
| **12** Univ Edinburgh | 129 | **37** Tsinghua Univ | 76 |
| **13** Northwestern Univ | 125 | **38** Univ Freiburg | 76 |
| **14** CALTECH | 122 | **39** Lawrence Berkeley Natl Lab | 75 |
| **15** Univ Minnesota | 120 | **40** Tianjin Univ | 75 |
| **16** Boston Univ | 119 | **41** Univ Calif Los Angeles | 73 |
| **17** Univ Oxford | 114 | **42** Yale Univ | 73 |
| **18** Univ Washington | 114 | **43** Univ Copenhagen | 70 |
| **19** Tech Univ Denmark | 109 | **44** UCL | 69 |
| **20** Univ Manchester | 107 | **45** Univ Warwick | 68 |
| **21** Univ Calif San Francisco | 98 | **46** Korea Adv Inst Sci & Technol | 67 |
| **22** Univ Penn | 97 | **47** Univ Maryland | 66 |
| **23** Duke Univ | 95 | **48** Arizona State Univ | 65 |
| **24** Univ Bristol | 93 | **49** Kyoto Univ | 64 |
| **25** Johns Hopkins Univ | 92 | **50** Newcastle Univ | 64 |

Source: Analysis of Web of Science publication records (2000 to mid-July 2018), Shapira et al. 2017 synthetic biology search strategy, N=11,369. Fuzzy list clean up (using organization name without department). UK organizations shaded in blue. \*Part year.

One half (25) of the top 50 institutions publishing in synthetic biology (by publication count) are based in the United States, led by MIT, Harvard, and the University of California, Berkeley. Nine of the top 50 institutions in this publication count list are in the UK, including Imperial College London, the University of Edinburgh, the University of Oxford, and the University of Manchester. Four institutions are in Japan, three in China, and two each in Switzerland and Denmark.

### International Co-authoring in Synthetic Biology

Co-authorship is the norm in synthetic biology, as in most other areas of science. Of the 11,368 synthetic biology publications in our 2000-2018* (*part year) WoS dataset, only 10% have a single author. Over one-third (34.1%) of the publications have 2-3 authors, while more than another third (35.4%) have 4-6 authors. Another 15.6% have 7-10 authors.

Co-authorship can occur among researchers in the same institution, in the same country, and across international boundaries. There are variations in the propensity of countries for international coauthorship. For example, among the four leading producers of synthetic biology publications (by publication counts), just over one quarter of US publications are internationally co-authored, with authors from China, the UK, and Canada being the three most prevalent. For the UK, 43% of synthetic biology publications are internationally-co-authored, most noticeably with the US (about 15% of all UK synthetic biology publications), followed by Germany and France. For the UK, 21% (271, non-duplicated) of its synthetic biology publications (2000-2018 part year) are with researchers from other member states of the European Union (EU). The four largest EU co-authoring partners for the UK are Germany (6.0%), France (4.3%), Italy (3.9%) and the Netherlands (3.3%). Germany also has a high international co-authorship rate (46% of all its synthetic biology publications), with the US, the UK, and Switzerland as the leading three international co-authors for German researchers. In China, international co-authorships comprise just over one-third of its synthetic biology publications, with the US contributing to nearly one fifth of Chinese publications in the domain. The UK is the second most common international co-authorship partner for Chinese researchers. (Table 3.)

**Table 3.** International Co-authoring, Top Four Synthetic Biology Publishing Countries, 2000-2018*.

| Country | Records | Internationally-authored<sup>+</sup> |  | Leading Co-Authoring Countries<sup>++</sup> |  |  |
| --- | --- | --- | --- | --- | --- | --- |
|  | Number | Number | Percent | 1<sup>st</sup> | 2<sup>nd</sup> | 3<sup>rd</sup> |
| **USA** | 4759 | 1242 | 26.1% | China 199 (4.2%) | UK 194 (4.1%) | Canada 140 (2.9%) |
| **UK** | 1290 | 559 | 43.3% | USA 194 (15.0%) | Germany 77 (6.0%) | France 55 (4.3%) |
| **China** | 1043 | 354 | 33.9% | USA 199 (19.1%) | UK 43 (4.1%) | Japan 29 (2.8%) |
| **Germany** | 1016 | 470 | 46.3% | USA 121 (11.9%) | UK 77 (7.6%) | Switzerland 58 (5.7%) |

Source: Analysis of Web of Science publication records (2000 to mid-July 2018), Shapira et al. 2017 synthetic biology search strategy, N=11,369. \*Part Year. +Non-duplicated. ++Co-authorships are for named country.<sup>8</sup> (Authors from multiple countries may contribute to some of these publications; hence totaling international co-authorships for each country with a subject country will exceed the total of subject country non-duplicated co-authorships).

A global mapping, for the top 15 countries by synthetic biology publications, demonstrates the dominant role of the US in synthetic biology international co-authorship linkages (Figure 3). The US leads as a synthetic biology co-authorship partner for the other top countries. The UK, China and Germany also serve as next tier hubs in international synthetic biology co-authorships. In addition to the dominant US network, there is also a European network (including the UK, Germany, France, Switzerland, Spain and Italy) and a Chinese network (involving the UK, US, Japan, Germany, and Australia among other countries).

**Figure 3.**
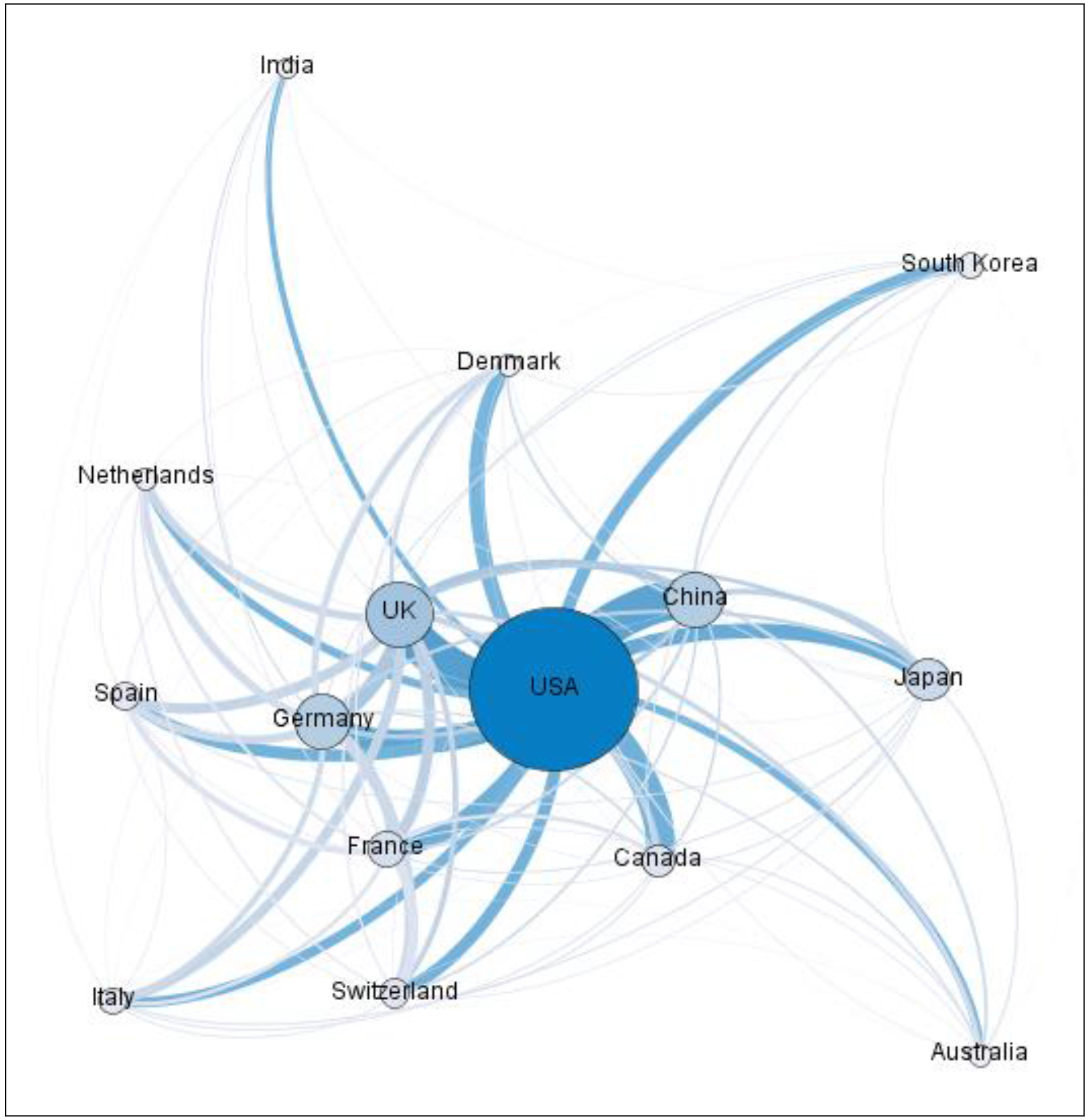
Synthetic Biology Co-Authorship Map, Top 15 Countries, 2000-2018*. Source: Analysis of Web of Science publication records (2000 to mid-July 2018), Shapira et al. 2017 synthetic biology search strategy, N=11,369. *Part Year. Top 15 countries, co-author network; circles are proportional to number of publications for respective country, edge size is proportional to the number of publications co-authored by researchers in those two countries.

### UK Co-Authoring in Synthetic Biology

Mapping co-authorship linkages of synthetic biology publications demonstrates the hub roles of the four leading locations by publication counts: Imperial College, the University of Edinburgh, the University of Oxford, and the University of Manchester. The University of Cambridge, Bristol University, the University of Warwick, and University College London also are among those with noticeable hub roles. (Figure 4.)

**Figure 4.**
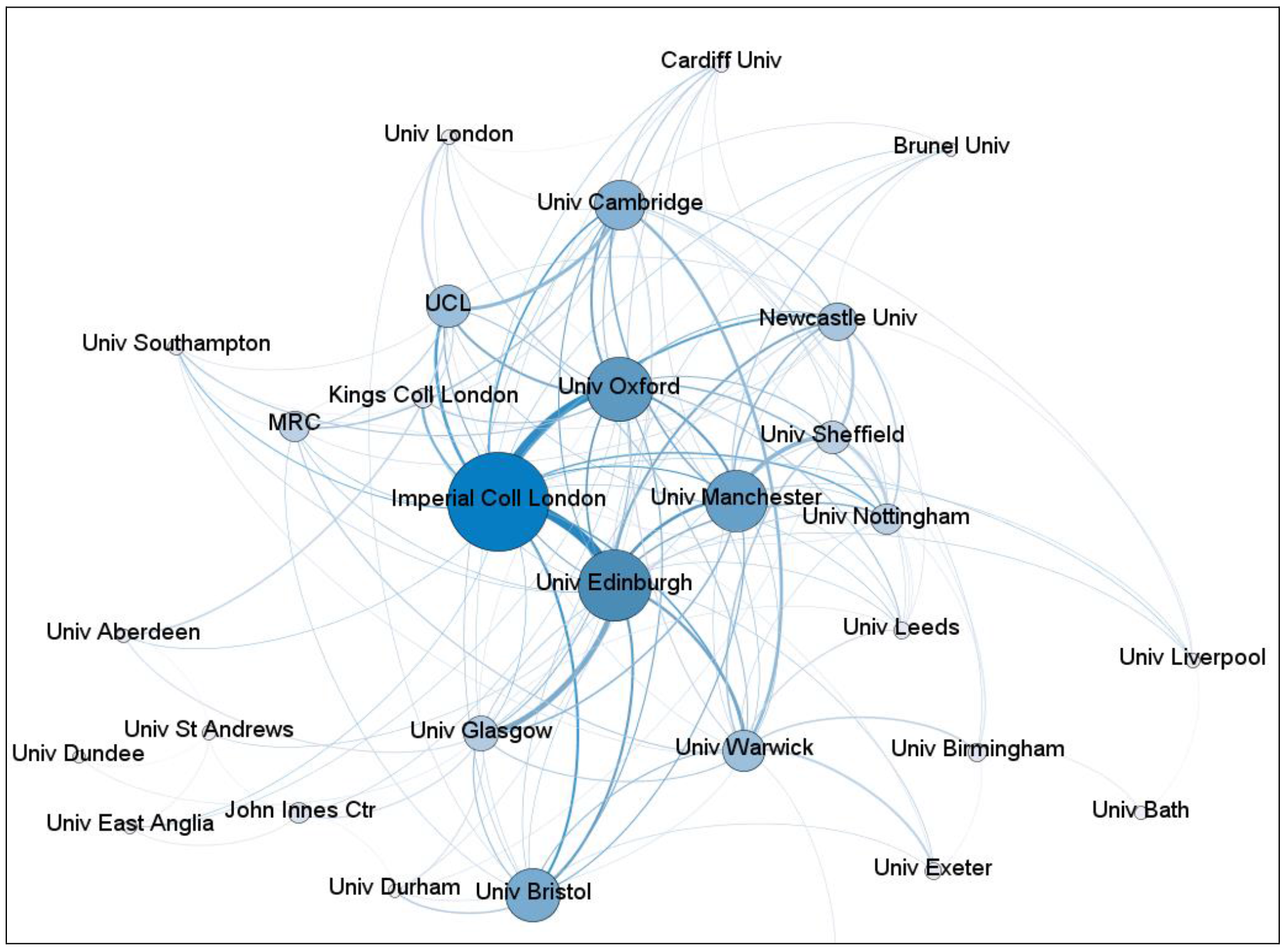
Synthetic Biology Publications, UK Co-Authorship Linkages. Source: Analysis of Web of Science publication records (2000 to mid-July 2018), Shapira et al. 2017 synthetic biology search strategy, N=1290. Top UK 30 organizations (by publication counts). Only co-authorship links among the UK organizations shown (i.e. other UK and international co-authorship links not shown). Circles are proportional to number of publications for respective organization; edge size is proportional to the number of publications co-authored by researchers in those two linked organizations.

### Leading Subject Categories Addressed by Synthetic Biology

The Web of Science allocates every journal, book or other publication that it records to at least one subject category. Currently, there are more than 250 WoS subject categories.^9^ These subject categories broadly relate to disciplines, although there are also some general and interdisciplinary categories. Synthetic biology is a crosscutting domain that covers multiple disciplines of science and technology and, indeed, seeks to engage and recombine key disciplines such as biology and biotechnology, engineering, and computing. Overall, our analysis indicates that synthetic biology publications encompass 188 or about three-quarters of WoS subject categories (Table 4).

**Table 4.** Leading Synthetic Biology Subject Categories.

|  |  | USA | UK | China | Germany | Worldwide |
| --- | --- | --- | --- | --- | --- | --- |
| 1 | Biochemistry & Molecular Biology | 22.2% | 19.1% | 14.3% | 21.8% | 20.7% |
| 2 | Biotechnology & Applied Microbiology | 17.2% | 13.6% | 32.1% | 21.4% | 20.2% |
| 3 | Biochemical Research Methods | 17.8% | 16.0% | 12.9% | 14.5% | 15.5% |
| 4 | Multidisciplinary Sciences | 12.8% | 14.7% | 11.4% | 10.9% | 11.2% |
| 5 | Chemistry, Multidisciplinary | 8.1% | 8.5% | 8.0% | 8.9% | 7.8% |
| 6 | Cell Biology | 5.8% | 4.0% | 3.2% | 5.6% | 4.8% |
| 7 | Microbiology | 4.0% | 5.5% | 5.2% | 5.4% | 4.8% |
| 8 | Biophysics | 3.9% | 3.3% | 4.1% | 5.0% | 4.4% |
| 9 | Genetics & Heredity | 4.2% | 2.6% | 2.5% | 2.8% | 3.7% |
| 10 | Mathematical & Computational Biology | 3.0% | 4.7% | 3.0% | 3.9% | 3.6% |
| 11 | Plant Sciences | 2.3% | 3.2% | 3.9% | 4.5% | 2.8% |
| 12 | Nanoscience & Nanotechnology | 2.3% | 2.6% | 4.0% | 2.1% | 2.6% |
| 13 | Medicine, Research & Experimental | 2.7% | 1.9% | 1.6% | 1.6% | 2.6% |
| 14 | Chemistry, Physical | 2.2% | 2.8% | 2.4% | 3.2% | 2.3% |
| 15 | Engineering, Electrical & Electronic | 2.2% | 3.0% | 1.3% | 0.4% | 2.3% |
| 16 | Biology | 2.0% | 2.9% | 2.3% | 2.2% | 2.3% |
| 17 | Materials Science, Multidisciplinary | 2.2% | 2.3% | 2.5% | 2.8% | 2.1% |
| 18 | Chemistry, Analytical | 1.6% | 1.6% | 3.5% | 1.1% | 2.0% |
| 19 | Chemistry, Medicinal | 1.8% | 1.2% | 1.2% | 2.3% | 1.9% |
| 20 | Pharmacology & Pharmacy | 1.3% | 1.2% | 0.9% | 1.9% | 1.8% |
| 21 | Engineering, Chemical | 1.4% | 0.5% | 2.4% | 0.8% | 1.6% |
| 22 | Immunology | 1.6% | 0.9% | 0.9% | 2.1% | 1.6% |
| 23 | Computer Science, Interdisciplinary Applications | 1.0% | 2.0% | 1.3% | 1.1% | 1.5% |
| 24 | Computer Science, Theory & Methods | 0.9% | 2.4% | 1.0% | 1.7% | 1.5% |
| 25 | Chemistry, Organic | 1.2% | 1.2% | 1.0% | 2.0% | 1.5% |
| 26 | Engineering, Biomedical | 1.6% | 1.6% | 1.2% | 0.6% | 1.5% |
| 27 | Computer Science, Artificial Intelligence | 0.7% | 2.2% | 1.1% | 1.3% | 1.3% |
| 28 | Physics, Applied | 0.9% | 1.2% | 1.1% | 1.5% | 1.2% |
| 29 | Automation & Control Systems | 1.1% | 1.8% | 0.8% | 0.5% | 1.0% |
| 30 | Materials Science, Biomaterials | 0.9% | 0.5% | 1.3% | 0.9% | 0.9% |

Source: Analysis of Web of Science publication records (2000 to mid-July 2018), Shapira et al. 2017 synthetic biology search strategy, N=11,369. Shows top 30 (by worldwide publication count) of 188 WoS Subject categories that include synthetic biology publications. A publication can be assigned more than one subject category, so totals of subject categories exceed 100%. \*Part Year.
<sup>9</sup> https://images.webofknowledge.com/images/help/WOS/hp_subject_category_terms_tasca.html

Worldwide, the top three WoS subject categories for synthetic biology publications are Biochemistry and Molecular Biology, Biotechnology & Applied Microbiology, and Biochemical Research Methods. For the first and third categories, UK synthetic biology publication outputs as a percentage of the total are similar to those observed worldwide, although the UK is a bit lower than worldwide level for Biotechnology & Applied Microbiology. Compared with the worldwide levels, China’s synthetic biology publication outputs are proportionately higher for Biotechnology & Applied Microbiology but lower in the other two top classifications. Both the US and the UK have above worldwide average levels of synthetic biology publications in the multidisciplinary subject classification. This includes prestigious multidisciplinary journals with exceptionally high impact factors such as *Nature* and *Science*, as well as recognized multidisciplinary journals such as *PloS One, Proceedings of the National Academy of Sciences (USA), Science Reports* (Nature), and *Nature Communications*. Among the next set of subject categories, and again compared with worldwide levels, the UK has a greater relative proportion of its synthetic biology publication outputs in Microbiology, Mathematical & Computational Biology, Plant Sciences, Engineering, Electrical & Electronic, Biology, and Chemistry, Physical, Computer Science, Theory & Methods, Computer Science, Artificial Intelligence, Computer Science, Interdisciplinary Applications, and Automation & Control Systems. The later set is of interest, indicating a relatively higher proportion of UK synthetic biology research activity related to information, data and automation technologies. (Table 4.)

To visualize the subject distributions and interdisciplinary linkages between subjects, we overlay our synthetic biology publication dataset onto a base map of science. ^10^ Our dataset covers synthetic biology publications in the WoS from 2000 through to mid-July 2018. While the visualization for all publications worldwide shows a spread of synthetic biology publications across disciplines, there are larger macro clusters in “Biochemistry, Molecular and Cell Biology,” “Biotechnology” (including plant sciences), and “Chemistry” (including biochemical research methods and chemistry, multidisciplinary). UK synthetic biology publications on the map of science are broadly comparable, with relatively more prominent clustering in multidisciplinary sciences and relatively greater linkages with computer science and engineering. (Figure 5.)

**Figure 5.**
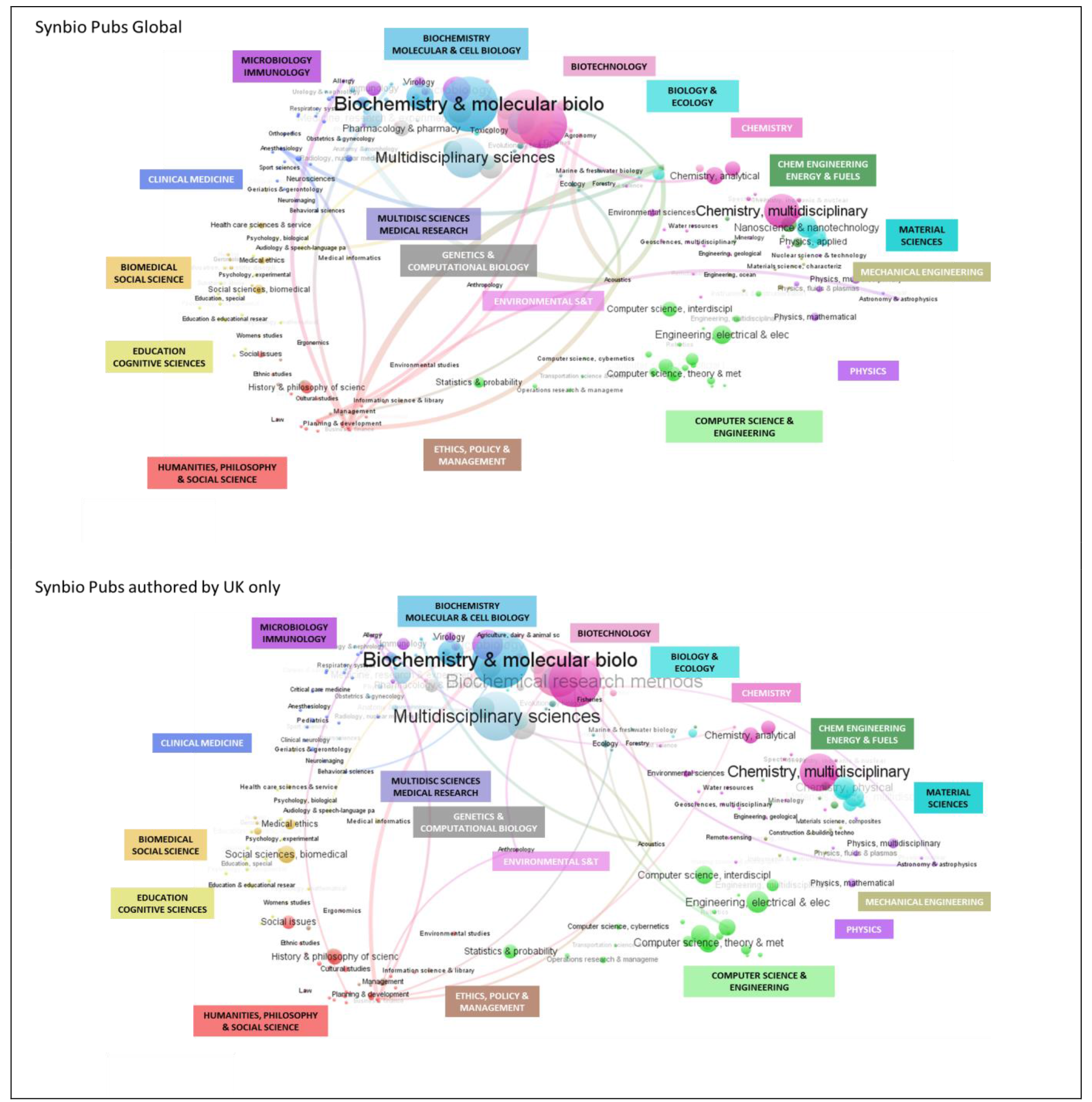
Synthetic Biology Publications on the Map of Science, Global and UK, 2000-2018*. Source: Analysis of Web of Science publication records (2000 to mid-July 2018), Shapira et al. 2017 synthetic biology search strategy, N=11,369. *Part Year. Map of science method from Carley et al. (2016), using VOSviewer (Van Eck and Waltman 2016), with customization of 2015 WoS 18-category macro-discipline labels for synthetic biology. Shows top 30 (by worldwide publication count) of 188 WoS Subject categories that include synthetic biology publications. A publication can be assigned more than one subject category, so totals of subject categories exceed 100%.

### Citations to Synthetic Biology Publications

Using citations to scientific publications as measures of research quality and impact is an imperfect but nonetheless commonly used indicator.^11^ Bearing in mind the limitations of using citation measures, we do observe significant differences among the top synthetic biology publishing countries in mean citations to papers attributed to one or more authors affiliated with institutions in those countries. Two North American countries – Canada and the US – both average more than 30 citations per publication in synthetic biology (noting that the US produces 10 times more publications than Canada). Switzerland averages almost 24 citations per publication, with Germany, the UK, and Netherlands each averaging around 20 citations per publication. Although third-ranked by publication count, China has the lowest mean citation rate among the top 15 synthetic biology countries. (Figure 6.)

**Figure 6.**
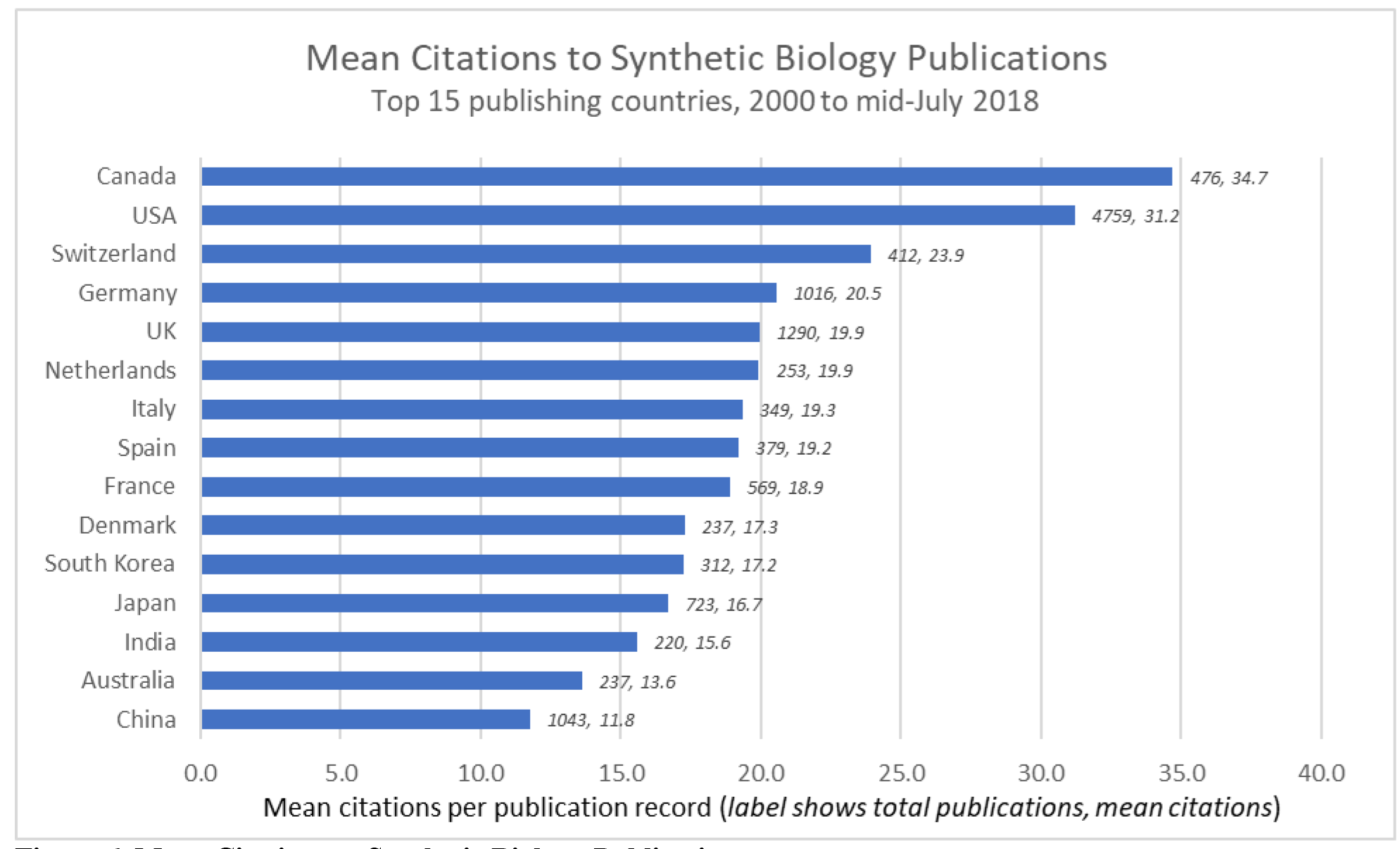
Mean Citations to Synthetic Biology Publications. Source: Analysis of Web of Science publication records (2000 to mid-July 2018), Shapira et al. 2017 synthetic biology search strategy, N=11,369.

### Funding Sponsorship

The top 15 funding organizations, by publication counts, in synthetic biology account for about 42% of all records in the synthetic biology publications data set.^12^ Among these top 15 funding organizations, six are in North America (with five in the US), five are in Europe (including two in the UK), and four are located in Asia (with two in China and Japan respectively). (Figure 7.) A publication may acknowledge funding from more than one sponsor, with some papers sponsored by funders in different countries. The highest mean citations per publication are garnered by publications that acknowledge funding from the US Office of Naval Research, followed by the US National Institutes of Health (NIH), the US Defense Advanced Research Projects Agency, the US Department of Energy (DOE), and the US National Science Foundation (NSF).

**Figure 7.**
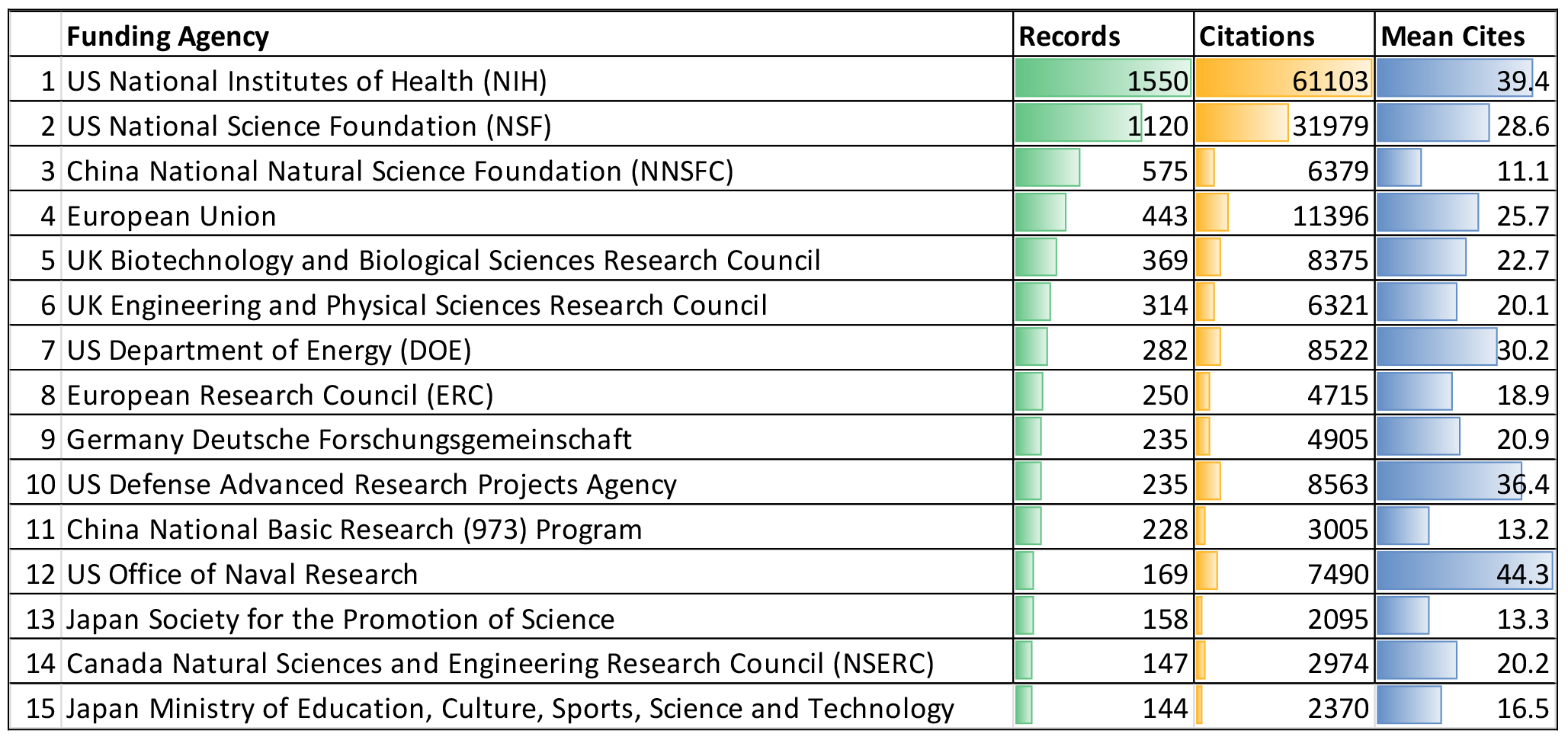
Citations to Publications sponsored by Top 15 Synthetic Biology Funding Agencies, 2000-2018*. Source: Analysis of Web of Science publication records (2000 to mid-July 2018), Shapira et al. 2017 synthetic biology search strategy, N=11,369 (of which 67% report funding acknowledgement information). VantagePoint used for list cleaning of funding agency organizational names.

## Synthetic Biology Patents

A granted patent provides exclusive rights to an invention and allows the patent owner to prevent others from using the patented invention for a period (typically twenty years). The broad policy objectives of patenting are to incentivize research and development of useful and novel applications by allowing patent owners time to recoup costs and generate revenues and to encourage disclosure to facilitate technological progress. An application for a patent does not mean that the relevant patent examining authority will grant the patent. Additionally, the grant of a patent does not necessarily mean that there will be use, commercialization, or licensing of the invention. In short, patenting is not the same as innovation, as there are multiple other steps and factors involved in the commercialization of new technology. However, from a technological landscape scanning perspective, patent applications do provide useful signals. Patenting activity can indicate interest in the exploitation of a new technology, particularly from a corporate perspective (as corporation are the leading filers of patents in most countries), and can suggest fields of invention that are viewed as promising. Patents can serve as signals for venture financing and can generate markets for inventions, although they can also be deployed in strategic ways to extract economic returns.

It is in the nature of emerging technologies that relevant patents are not always easily defined, especially as standard patent classifications typically lag in being updated and also because patent applicants may use new terminology or deliberately not use specific terminologies. We have developed a method for discerning synthetic biology patents based on identifying a core set of relevant patents and keywords, then extending from this corpus to other relevant patents using a citation-tree algorithm (see section on methods and sources for added details and references). For the priority year period 2003 to August 3, 2018, from 9,263 Derwent Innovations synthetic biology patent records we matched 8,460 PATSTAT basic patent application records (for the original invention in a patent family) to obtain assignee geographical information. There are over 10,600 original patent assignees named in these records, comprised of about 3,600 original assignees standardized by name in PATSTAT. Nearly 16% of the patent records indicate more than one assignee. Some 36% of the assignees are associated with multiple patents, while some assignees are part of larger multinational corporations. The majority of patents are associated with multiple inventors (for patents where inventor names are recorded, just 5% are single inventor patents). In total, our synthetic biology patent records name about 23,000 inventors, of whom 23% are inventors who have filed more than one patent.

As in many patent landscapes searches, we acknowledge that the boundaries of the domain, in this case synthetic biology, are hard to delineate. We have pursued a broad definition of the domain (noting that few patents actually use the exact terminology “synthetic biology”). We recognize that there are inevitably trade-offs in precision and recall. At an aggregated level, however, the search approach is useful for indicating broad patenting trends.

We primarily analyze patent applications. Unless otherwise indicated, the use of the term “patent” in the text denotes a patent application. In addition to the caveats noted above (e.g. patent applications may signal interests in innovation but they are not in themselves innovations), patent applications are usually not disclosed until at least 18 months from being filed, and there are procedures that applicants can use to further extend disclosure. This means that patent application records for the most recent years are not complete.

### Growth in Synthetic Biology Patenting

Our analysis indicates upward worldwide growth in synthetic biology patent applications. Synthetic biology publications also have an upward worldwide growth (Figure 8). From 2007 through to 2011, there was a noticeable increase in synthetic biology patent applications. Since 2011, the growth rate of patenting tracks that of publishing, suggesting broadly comparable growth levels of global activity in knowledge exploration and invention exploitation in the synthetic biology domain. (We again note that the apparent drop-off in published patent applications from 2015-2016 to the present should not be taken as a real decline, as it reflects the gap between patent filing and publication. This gap is typically 18 months but can be longer in some circumstances.)

**Figure 8.**
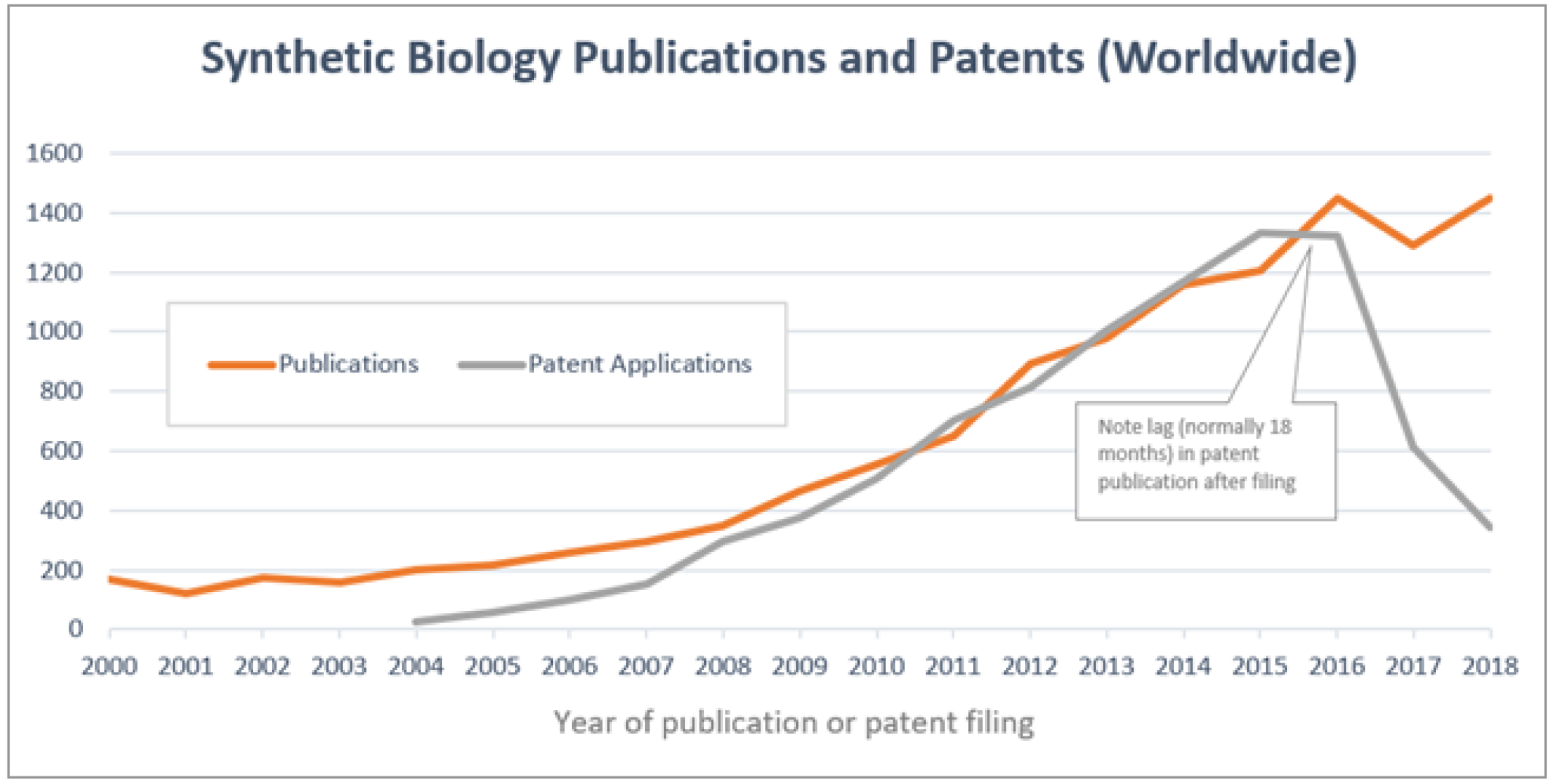
Synthetic Biology Publications and Patents. Source: Publications – analysis of Web of Science publication records (2000 to mid-July 2018), Shapira et al. 2017 synthetic biology search strategy, N=11,369. Patents – analysis of PATSTAT patent records (2003 to August 3, 2018), Kwon et al. 2016 synthetic biology patent search strategy, N=8,460. VantagePoint used for data cleaning and analysis.

The United States accounts for more than one-half of all synthetic biology patent applications, by location of the original patent assignee. (Figure 9). Assignees based in Japan account for about 9% of worldwide patents, followed by those located in Switzerland (5.5%, Germany (5%) and other European countries. Great Britain accounts for about 3% of worldwide patents. Our data does not as yet show a high level of Chinese synthetic biology patenting (there may be time lags in capturing applications from China’s State Intellectual Property Office).

**Figure 9.**
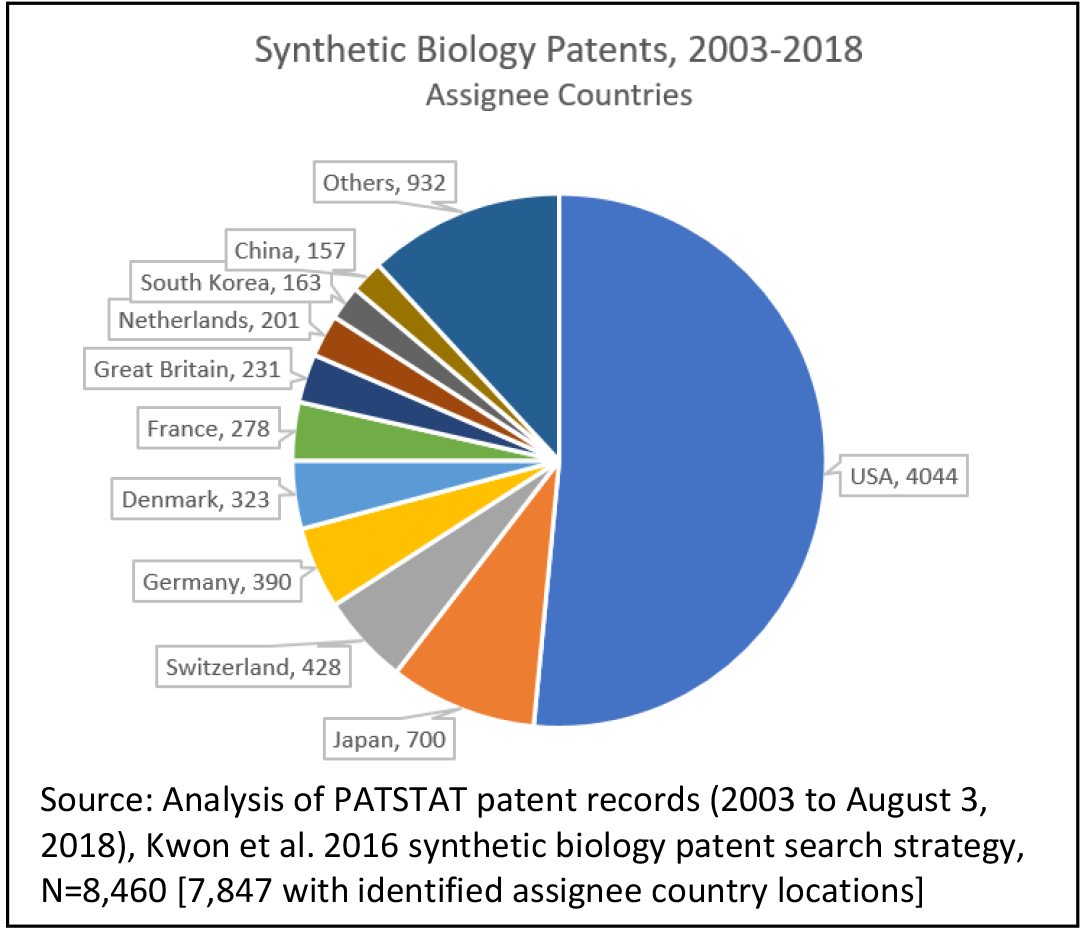
Synthetic Biology Patents, By Country.

### National Patenting Variations

While at an aggregated global scale, the growth of synthetic biology publications has been mirrored by comparable growth in synthetic biology patent applications, there are significant variations among the leading countries. Denmark, Switzerland and Japan have higher rates of synthetic biology patenting relative to their synthetic biology publication authorships. The USA and the Netherlands patent at rates only slighly below their level of publications. Synthetic biology patenting by assignees located in Great Britain is low (.20) relative to the high rate of synthetic biology publishing by UK researchers.

**Figure 10.**
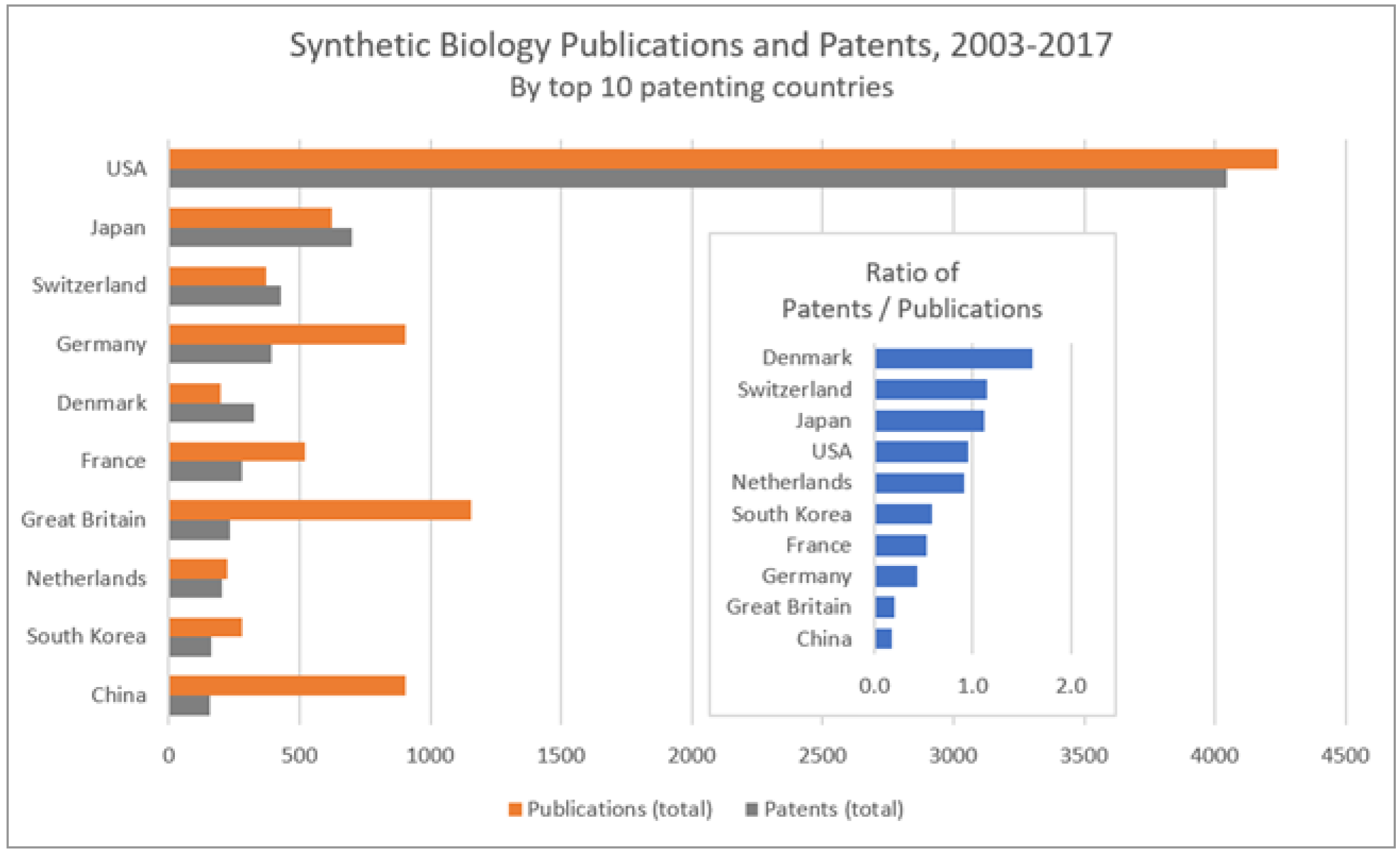
Synthetic Biology Publications and Patents, 2003-2017. Source: Publications – analysis of Web of Science publication records (2000 to mid-July 2018), Shapira et al. 2017 synthetic biology search strategy, N=11,369. Patents – analysis of PATSTAT patent records (2003 to August 3, 2018), Kwon et al. 2016 synthetic biology patent search strategy, N=8,460. VantagePoint used for data cleaning and analysis.

### Leading Patent Assignees

Companies are most frequent among the world’s leading patent assignees in the synthetic biology domain, with 27 companies present among the top 41 assignees. This top group also contains universities and non-profit research organizations, particularly in the US and to a lesser extent in Japan, France, and Singapore. The US is the leading location for patent assignees, with 26 organizations, followed by Denmark and Japan (each with three) and France and Switzerland (two each). (Figure 11.) The leading British patent assignee is the Glaxo Group / GlaxoSmithKline (76^th^ globally). There are six companies among the top (11) British patent assignees, four universities and one governmental non-profit research organization. (Figure 12). When reviewing these results, keep in mind our previous caveats about limitations and trade-offs in precision and recall in identifying synthetic biology patents.

**Figure 11.**
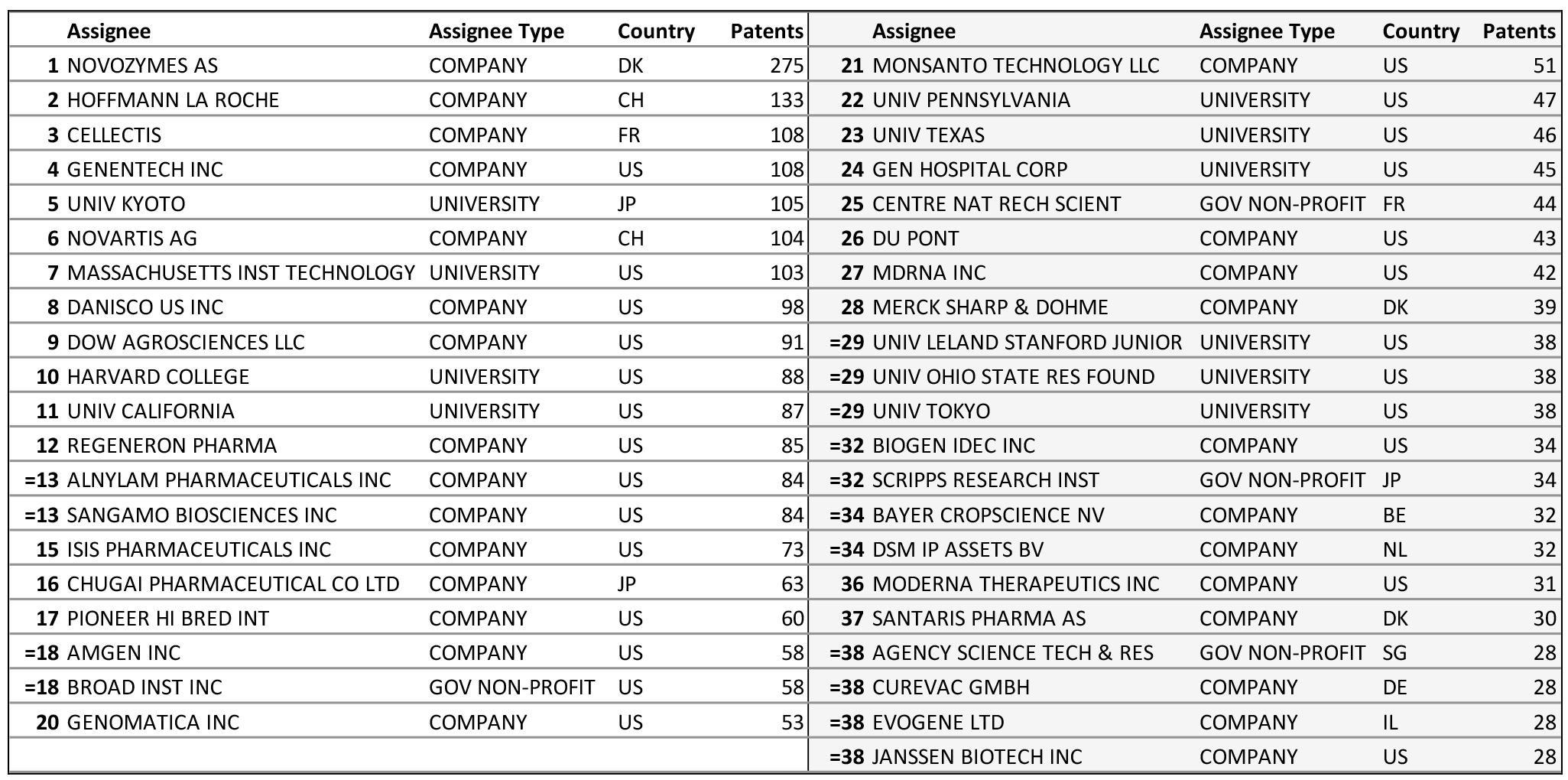
Top Patent Assignees in the Synthetic Biology Domain, Worldwide. Analysis of Patstat patent records (2003 to August 3, 2018), Kwon et al. 2016 synthetic biology patent search strategy, N=8,460 [7,847 with identified assignee country locations]

**Figure 12.**
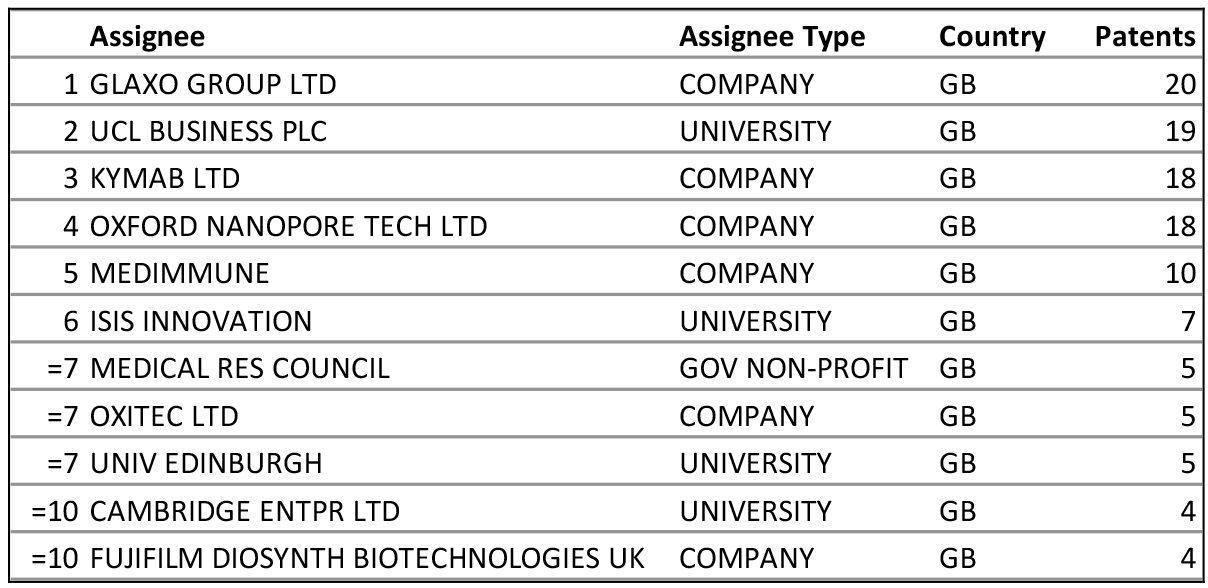
Top Patent Assignees in the Synthetic Biology Domain, Great Britain. Analysis of PATSTAT patent records (2003 to August 3, 2018), Kwon et al. 2016 synthetic biology patent search strategy, N=8,460 [7,847 with identified assignee country locations]

### Synthetic Biology Patenting Visualized on the Map of Patents

Synthetic biology patenting covers a broad range of technological and application areas. Just over 100 International Patent Classification (IPC) subclasses (or about 16% of all IPC subclasses) are represented in our synthetic biology patent data set, with the largest patent subclasses including mutation or genetic engineering (C12N), measuring or testing processes involving enzymes, nucleic acids or microorganisms (C12Q), and medicinal preparations containing genetic material (A61K). An individual patent may be classed under multiple IPCs. To visualize the distribution of synthetic biology patents, we have mapped the worldwide synthetic biology patent set onto a base map of patents, organized into 35 major technological groups (Figure 13). This highlights major clusters in areas of biotechnology, drugs, instrumentation, pesticides, and catalysts, with smaller groups in food, detergents, information technology, and filtration and fibers. A mapping of patents with assignees located in Great Britain is shown, on the same base map of patents, is also shown (Figure 14).

**Figure 13.**
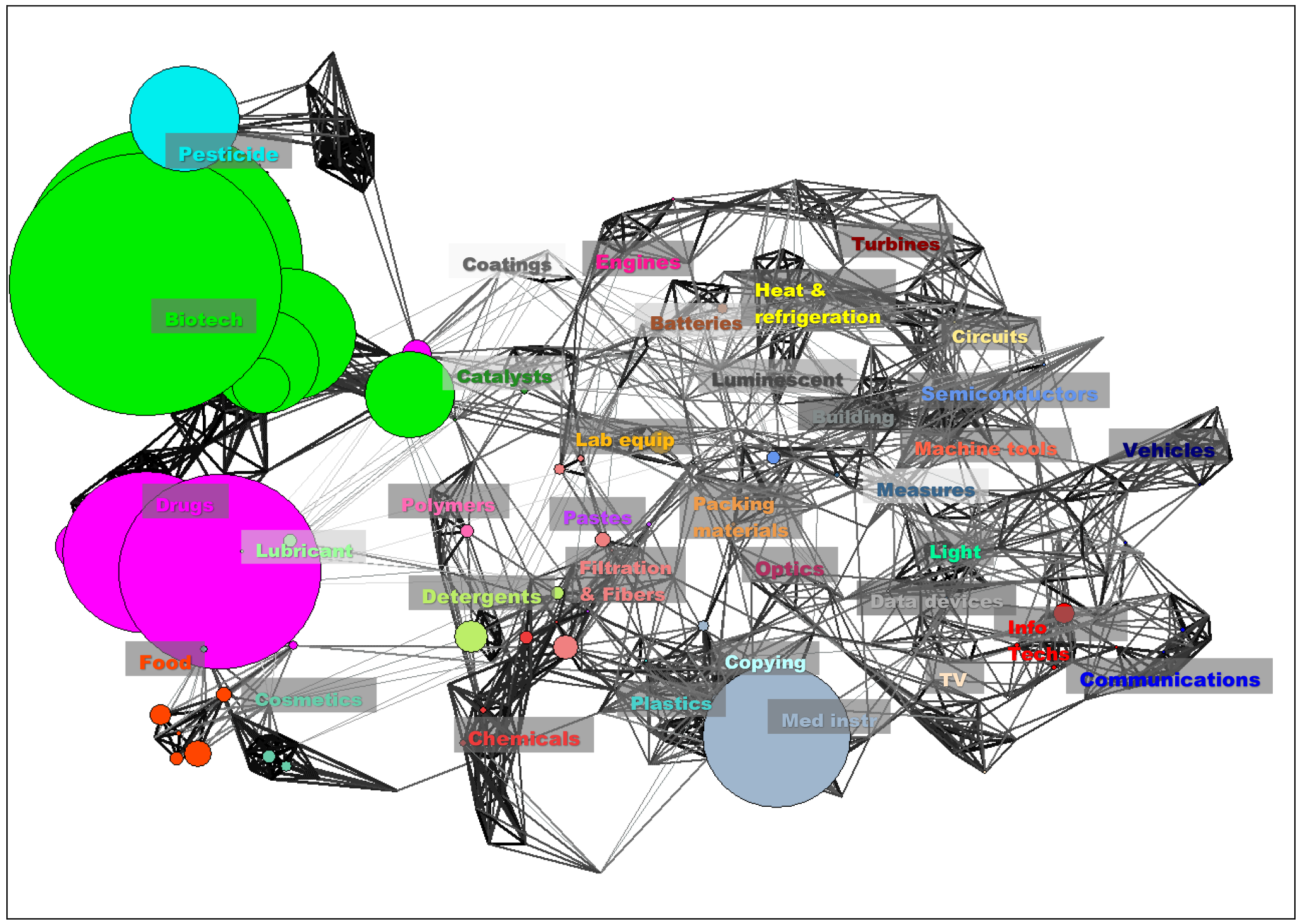
Synthetic Biology Patents on Map of Patents, Worldwide, 2003-2018. Source: Analysis of PATSTAT patent records (2003 to August 3, 2018), Kwon et al. 2016 synthetic biology patent search strategy, N=8,460. Overlaid on visual map of patents comprising 466 technological classifications and 35 technological groups (see Kay et al. 2014).^13^ PATSTAT IPC classifications. The size of nodes is proportional to the number of patent applications in the corresponding technology group. On the base map, lines represent relationships between technological categories (the darker the line, the shorter the technological distance between categories).

**Figure 14.**
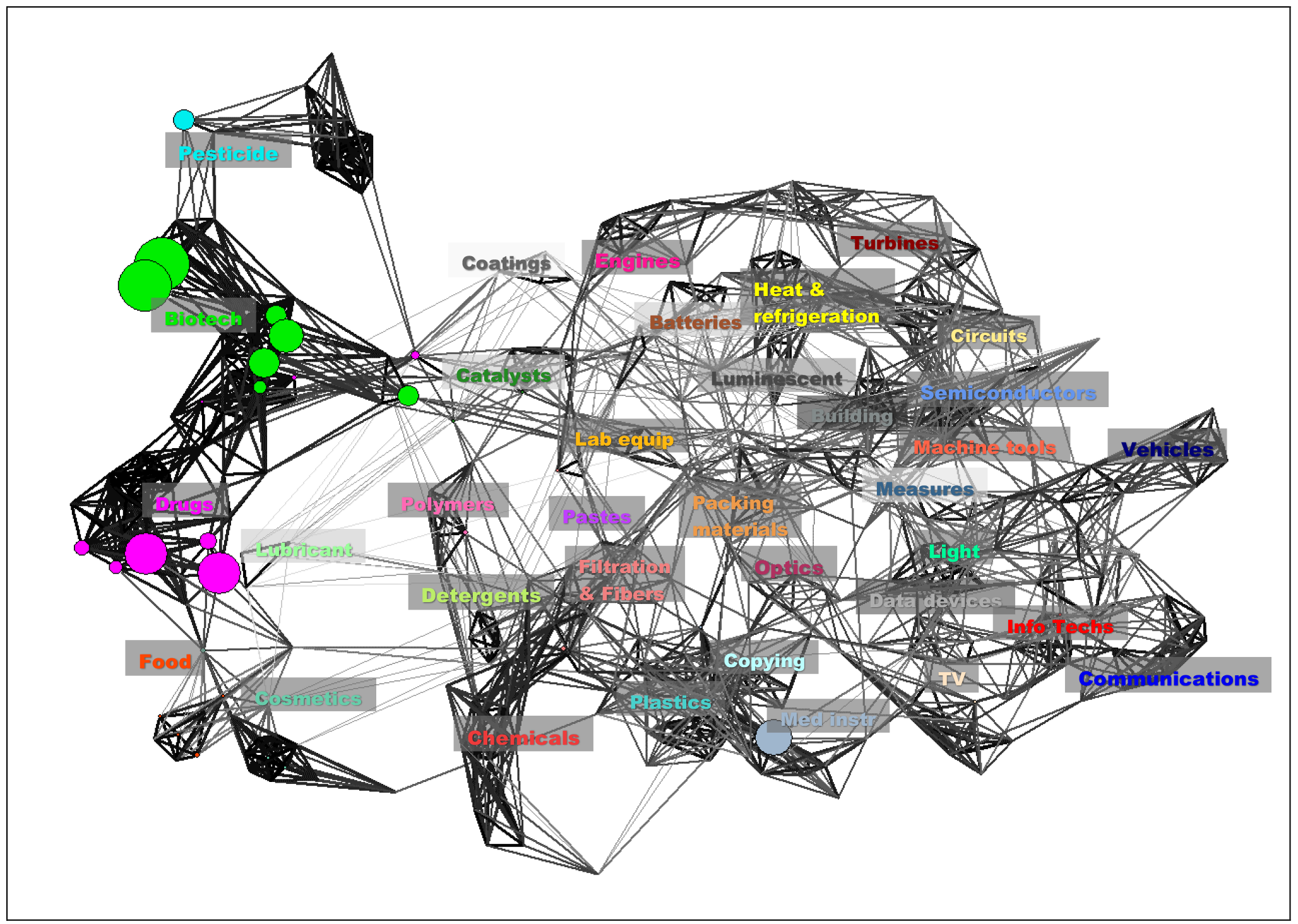
Synthetic Biology Patents on Map of Patents, Great Britain, 2003-2018. Source: Analysis of PATSTAT patent records (2003 to August 3, 2018), Kwon et al. 2016 synthetic biology patent search strategy, N=231 (assignees in Great Britain). Overlaid on visual map of patents comprising 466 technological classifications and 35 technological groups (see Kay et al. 2014). PATSTAT IPC classifications. The size of nodes is proportional to the number of patent applications in the corresponding technology group. On the base map, lines represent relationships between technological categories (the darker the line, the shorter the technological distance
between categories).

## Appendix

This appendix compares publication counts returned using the Shapira et al 2017 “expanded” search with a simple topic search for “synthetic biology.” A topic search captures the use of a term in the title, abstract, or key words or a publication. While broadly these searches track one another in terms of growth rates, the expanded search captures publications in the domain even where the topic term “synthetic biology” is not used. A third search for the use of “engineering biology” as a topic returns relatively few publications.

**Figure A1.**
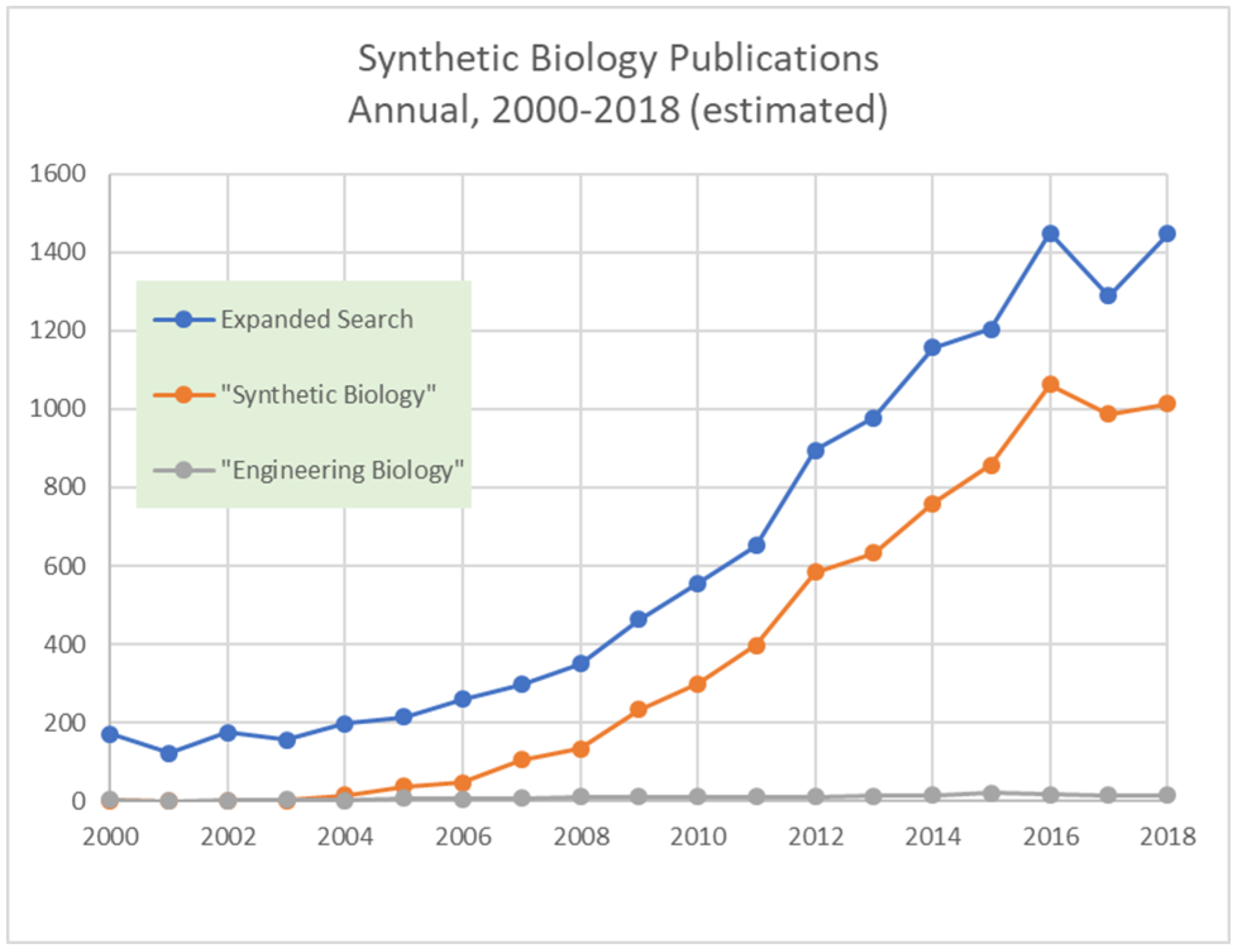
Worldwide Synthetic Biology Publications. Source: Analysis of Web of Science (WoS) publication records. Expanded search from Shapira et al. 2017 synthetic biology search strategy, 2000 to mid-July 2018, N=11,369. WoS topic searches (title, abstract, or key words containing exact term), 2000 to mid-October 2018 for “Synthetic Biology” N= 7,000, and “Engineering Biology” N=173. Annualized totals for 2018 estimated from part-year 2018 publication trends. The WoS databases searched comprised SCI-EXPANDED, SSCI, CPCI-S, CPCI-SSH, and A&HCI (document type: all).

| Added statistics | Publication counts (actual), 2000-2018*<br>(*see dates in source notes) |
| --- | --- |
| Expanded search | 11,369 |
| “Synthetic Biology” | 7,000 |
| “Engineering Biology” | 173 |
| “Synthetic Biology” OR “Engineering Biology” | 7,142 |
| “Synthetic Biology” AND “Engineering Biology” | 31 |

### Acknowledgements

The authors acknowledge support from the Biotechnology and Biological Sciences Research Council [grant number BB/M017702/1] (Manchester Synthetic Biology Research Centre for Fine and Speciality Chemicals) and the National Science Foundation [Award No. 1759960]. Findings and observations contained in this paper are those of the authors and do not necessarily reflect the views of the Biotechnology and Biological Sciences Research Council or the National Science Foundation.

## Footnotes

1 Clarivate Analytics. https://clarivate.com/products/web-of-science/databases/

2 Shapira P, Kwon S, Youtie J. Tracking the Emergence of Synthetic Biology, *Scientometrics*, 2017, 112: 1439-1469. http://dx.doi.org/10.1007/s11192-017-2452-5.

3 The WoS databases searched comprised SCI-EXPANDED, SSCI, CPCI-S, CPCI-SSH, and A&HCI (document type: all). This search updates the original search in Shapira et al. (2017), which covered the period 2000-2015.

4 https://www.thevantagepoint.com/

5 Clarivate Analytics. https://clarivate.com/products/derwent-innovation/

6 Kwon S, Youtie J, Shapira P. 2016. Building a Patent Search Strategy for Synthetic Biology. Working Paper. Georgia Tech Program in Science, Technology and Innovation Policy, Atlanta, GA, USA. March 10. http://bit.ly/2E3Py7T

7 European Patent Office. https://www.epo.org/searching-for-patents/business/patstat.html

8 Authors from multiple countries may contribute to some of the publications co-authored with a named subject country. Hence totaling international co-authorships for each country with a subject country will exceed the total of subject country non-duplicated co-authorships.

9 https://images.webofknowledge.com/images/help/WOS/hpsubjectcategorytermstasca.html

10 The base map represents the foundational organization of science, drawing on co-citation patterns for subject categories of WoS journals. See Porter, A. L., & Rafols, I. (2009). Is science becoming more interdisciplinary? Measuring and mapping six research fields over time. *Scientometrics, 81*, 719-745, doi: 10.1007/s11192-008-2197-2; and Rafols, I., Porter, A. L., & Leydesdorff, L. (2010). Science overlay maps: A new tool for research policy and library management. *Journal of the Association for Information Science and Technology*, 61(9), 1871-1897, doi: 10.1002/asi.21368. As noted in Shapira et al, 2017, “the map visualizes the distance and intensity of co-citations between corresponding subject categories. Each node represents individual WoS subject categories, with colors used to depict subject categories within the same macro-discipline group. The subject categories that are assigned to synthetic biology articles in our publication dataset are matched to the 18 macro-disciplines and displayed on the base map” and “the node size is proportional to the number of articles in that subject category.” For the method to construct the 18 macro disciplines, see Carley, S., Porter, A. L., Rafols, I., & Leydesdorff, L. (2017). Visualization of disciplinary profiles: Enhanced science overlay maps. *Journal of Data and Information Science*, 2(3), 68-111, https://content.sciendo.com/view/iournals/idis/2/3/article-p68.xml. The VOSviewer clustering method and algorithm is used, see: van Eck, N. J., & Waltman, L. (2010). Software survey: VOSviewer, a computer program for bibliometric mapping. *Scientometrics*, 84(2), 523-538. doi: 10.1007/s11192-009-0146-3.

11 On problems associated with using citation analyses, see, for example, MacRoberts, M. H. and MacRoberts, B. R. (1989), Problems of citation analysis: A critical review. Journal of the American Society for Information Science, 40: 342-349. doi:10.1002/(SICI)1097-4571(198909)40:5<342::AID-ASI7>3.0.CO;2-U.

12 67% of synthetic biology publication records report funding acknowledgments information.

13 Kay, L., Newman, N., Youtie, J., Porter, A. L. and Rafols, I. (2014), Patent Overlay Mapping: Visualizing Technological Distance. *J Assn Inf Sci Tec*, 65: 2432-2443. doi: 10.1002/asi.23146

